# Sharing the burden? Earthworms and woodlice show seasonal complementarity in peak abundances in soil in an oak-beech temperate woodland

**DOI:** 10.1101/2020.11.13.381863

**Authors:** Daniel Carpenter, Emma Sherlock, Kelly Inward, Kerry Carroway, Angela Lidgett, Paul Eggleton

**Affiliations:** Soil Biodiversity Group, Department of Life Sciences, Natural History Museum, London, SW7 5BD, UK; Aquatic Invertebrates Division, Department of Life Sciences, Natural History Museum, London, SW7 5BD, UK

**Keywords:** decomposers, earthworms, complementarity, millipedes, seasonal patterns, woodlice

## Abstract

Complementarity between functional analogues can confer resistance and resilience on ecosystems in the face of environmental change. High biodiversity can lead to increased ecosystem functionality through complementary effects. Earthworms, woodlice and millipedes can have high densities in leaf litter and soils, but little is known about their seasonal patterns. The two groups play important roles in the breakdown and incorporation of organic matter into soils. Differences in peak abundance could affect the rates of litter break down and incorporation in different seasons. We sampled earthworms, woodlice and millipedes from leaf litter soil every month for ten years in a New Forest woodland. We used non-parametric regression to explore monthly and yearly variation in the abundance of decomposer organisms and soil temperature and moisture. Earthworms have a distinct seasonal peak in density different from woodlice and millipedes. Earthworm peak density is in the winter and spring and is correlated with greatest soil moisture. Woodlice (and millipede) have their peak density is in the summer and is correlated with the highest soil temperatures. This means that earthworms, woodlice and millipedes have complementary peaks in abundance. These two groups have similar functional roles in litter decomposition and these data imply ecological complementarity in this important ecological process. This effect is likely to be widespread in lowland woodland in the UK and Europe, with only extreme temperatures and low pH limiting the distribution. Increased summer drought as a result of climate change may lead to changes in the relative abundance of these three groups and in particular local extinctions of earthworms which will in turn affect litter decomposition.

## 1. Introduction

A global decline in biodiversity, as a result of both climatic and human-influenced environmental change, is fuelling a renewed focus on our understanding of the relationships between species diversity and ecosystem function. Specifically, the resilience of an ecosystem to change depends on its ability to maintain function while species diversity declines (Yachi and Loreau, 1999). Ecological resilience therefore depends on the relative levels of functional redundancy and complementarity across species within an ecosystem.

Ecosystems are said to have functional redundancy when there are many species doing the same or similar jobs (Loreau, 2004). This implies that local extinction events could cause species to be lost from the system with no apparent loss in function, as other species compensate by supplying the function lost through the extinct species (e.g. Elser et al., 1995; Goheen et al., 2005). Functional redundancy implies ecological analogy, where similar species have essentially the same functional role.

Competition for resources between species, however, works against functional redundancy as species usually must evolve different strategies to exploit the same resource. This means that species tend to reach abundance equilibria based on the different strategies that they employ to exploit a common resource. These strategies are often where species use a different part of a common resource (Wilson et al., 1999) or exploit the resource at different temporal or spatial scales (Albrecht and Gotelli, 2001). In extensive heterogeneous environments (e.g. soil) with a large number of generalist organisms (e.g. decomposers), competition is generally reduced as species can avoid each other or use different parts of the same resource. This allows coexistence between species with similar environmental tolerances, which generally leads to high functional complementarity (Setälä et al., 2005).

Complementary effects can enhance ecosystem functioning through an increase in biodiversity (Loreau and Hector, 2001; Bracken and Stachowicz, 2006; Yachi and Loreau, 2006); process rates are often greater with higher biodiversity than with lower biodiversity. Complementarity also occurs when the combined function of two or more species can provide enhanced mutualistic benefits to other species, for example by reducing stress (Stachowicz and Whitlatch, 2005).

Decomposition is a vital biological process that breaks down dead organic matter, incorporates it into the soil and releases plant nutrients. The process is conventionally attributed predominantly to microbes, but it is increasingly obvious that macrofauna have an important role (Wall et al., 2008; Gessner et al., 2010.).

Invertebrates in the decomposer guild have four main roles in the decomposition process. First they increase the surface area of leaf litter by communition (Anderson, 1988). Second, they are direct decomposers of substrates through (mostly) midgut enzyme production (Hassall et al., 1987; Guggenberger et al., 1996; Le Bayon et al., 2002). Third, they act as habitats for microbial decomposition (Hassall et al., 1987; Pokarzhevskii et al., 1997; Jegou et al., 2001; Scullion et al., 2002; Drake and Horn, 2007). Fourth, many of the invertebrates are coprophages (Ullrich et al., 1992).

While much is known about the functional roles of these soil macrofauna groups, there is less information concerning their seasonal patterns (Gerard, 1967; Baker et al., 1993; Spurgeon and Hopkin, 1999; Tondoh, 2006). In temperate systems, seasonal differences in soil temperature and soil moisture affect the abundance of decomposer organisms (Riutta et al., 2012.), with peaks of abundance in a particular season as a result of their environmental tolerances. Previous work has shown the response of earthworms to seasonal and yearly changes in soil temperature and moisture over a six year period (Eggleton et al., 2009). Decomposition rates may well vary with season, due to differences in peak decomposer abundance, rates being high with peak abundance and low when abundance is low. If the peak abundance for the different decomposer groups coincides in a season, then rates of decomposition are likely to be greatest in that season and concomitantly low at times when abundances are low. Alternatively, if the peak abundance of decomposer groups are complementary (i.e. one group has its peak when another has low abundance) then decomposition rates may well be maintained throughout the year. However, partitioning of the leaf litter resource could also account for differences in seasonal abundances of different litter feeding taxa.

Here, we present data from a ten-year study of soil and leaf litter macrofauna from a broadleaf wood in southern England. We explore the temporal patterns of the three main decomposer taxa (earthworms, woodlice and millipedes) to uncover in which season each taxon has its peak abundance and to estimate the extent to which these peaks coincide or are complementary. We found that peak seasonal abundance for earthworms was different to that of woodlice and millipedes.

## 2. Materials and Methods

### 2.1 Site description and sampling methods

Whitley Wood is an ancient semi-natural wood situated in the New Forest, Hampshire, UK (Coordinates 50.849152, −1.5766458). It is classified broadly as a W10 *Quercus robur - Pteridium aquilinum - Rubus fruticosus* woodland under the National Vegetation Classification (NVC) (Rodwell, 1991). However, it has two co-dominant tree species, oak (*Quercus robur*) and beech (*Fagus sylvatica*), and a varied understorey of hawthorn (*Crataegus monogyna*) and field maple (*Acer campestre*).

The ground flora is patchy, but includes bracken (*Pteridium aquilinum*), bramble (*Rubus fruticosus agg*), common dog violet (*Viola riviniana*) and wood spurge (*Euphorbia amygdaloides*). The wood, like many of those in the New Forest, is browsed by ponies (Tubbs, 2001). Soils at the study site are Planosols (FAO WRB) and are seasonally wet, in places gleyed, indicating temporary waterlogging.

We sampled every month for ten years, from March 2002 to February 2012 in a 2 ha area of the wood. Each month we placed a 100m transect line at random within the 2 ha area and sampled at fifteen points at 7 m intervals along the transect. At each sampling point we used two methods to extract invertebrates from the leaf litter and soil respectively. To extract invertebrates from leaf litter we placed sieved leaf litter from a 1 m^2^ quadrat into Winkler bags for three days (Krell et al., 2005). To sample earthworms in the soil we dug four pits (15 cm × 15 cm × 10 cm [depth]) each next to the middle of the four edges of the 1 m^2^ quadrat that was used to extract leaf litter, giving a total of 60 pits per transect. Adult earthworms, woodlice and millipedes from these samples were identified to species.

We pooled data from each part of the transect (i.e. litter or soil separately) to give a single species × abundance list for each transect (as in Eggleton et al., 2009). Our sampling methods risk double counting all groups as the leaf litter scrapes inevitably samples some soil and the soil pits will sample some leaf litter. We have employed the simplifying assumption that the soil pits sample the earthworms almost comprehensively (as even epigeic earthworms are found predominantly in the soil and humus layers) and that sieving litter samples the woodlice and millipedes almost comprehensively (as those two groups are generally leaf litter inhabitants only).

At each sampling point we also measured soil moisture using a using a SM200 soil moisture probe attached to a HH2 meter (Delta T Devices, Cambridge, UK) and soil temperature using a simple bi-metal thermometer.

### 2.2 Data analysis

We partitioned the variance in the abundances of the species into seasonal and year components using non-parametric regression in the R package *sm* (Bowman and Azzalini, 1997, as outlined in Eggleton et al., 2009 (as modified from Fergusson et al., 2007). Full details are in the supporting information of Eggleton et al., 2009). Initial bandwidth values for the partitioning analysis were obtained using *sm.regression.autocor* as monthly points are likely to be autocorrelated.

Relationships between (log +1)-transformed abundances and soil moisture and soil temperature were also examined using non-parametric regression. In all cases we used the *sm.regression.autocor* function to estimate the optimum bandwidths for smoothing and the used the *sm.regression* function to test for the presence of a significant relationship, using the *model=”no effect”* command. Where the null hypothesis was rejected at the p<0.05 level an additional test was undertaken to examine whether the relationship could be modelled as linear, using the *model=”linear”* command.

## 3. Results

### 3.1 Decomposer data

Over the ten years of the project we sampled 5,279 adult earthworms in 15 species, 15,156 adult woodlice in five species and 438 adult millipedes in 11 species (see Table 1 for details). The five most abundant species of earthworm account for nearly 98% of the total earthworm abundance. For one of the five species of woodlice identified (*Haplophthalmus danicus* Budde-Lund, 1880), we only sampled one individual in the 120 sampling periods. The five most abundant millipede species account for nearly 93% of the total millipede abundance.

**Table 1.** Numbers of specimens counted for each species. Percentages are calculated as totals within each major clade. The woodlice and millipede data are all from litter samples and the earthworm data are all from soil pits.

| Species | No. sampled |
| --- | --- |
| Earthworms |  |
| *Dendrobaena octaedra* | 2679 (50%) |
| *Allolobophora chlorotica* | 1441 (27%) |
| *Dendrobaena attemsi* | 458 (9%) |
| *Aporrectodea caliginosa* | 373 (7%) |
| *Lumbricus rubellus* | 290 (5%) |
| *Aporrectodea rosea* | 71 (1%) |
| *Satchellius mammalis* | 10 (<1%) |
| *Octolasion lacteum* | 9 (<1%) |
| *Octolasion cyaneum* | 7 (<1%) |
| *Dendrodrilus rubidus* | 5 (<1%) |
| *Aporrectodea longa* | 2 (<1%) |
| *Dendrobaena hortensis* | 2 (<1%) |
| *Dendrobaena pygmaea* | 2 (<1%) |
| *Aporrectodea limicola* | 1 (<1%) |
| *Eiseniella tetraedra* | 1 (<1%) |
| Woodlice |  |
| *Philoscia muscorum* | 10013 (61%) |
| *Oniscus asellus* | 2336 (14%) |
| *Trichoniscus pusillus* | 1487 (9%) |
| *Porcellio scaber* | 1319 (8%) |
| *Haplophthalmus danicus* | 1 (<1%) |
| Millipedes |  |
| *Glomeris marginata* | 159 (36%) |
| *Polydesmus denticulatus* | 90 (21%) |
| *Chordeuma sp* | 74 (17%) |
| *Cylindroiulus punctatus* | 48 (11%) |
| *Polydesmus sp* | 35 (8%) |
| *Polydesmus coriaceus* | 18 (4%) |
| *Chordeuma proximum* | 10 (2%) |
| *Polyxenus lagurus* | 1 (<1%) |
| *Ophiulus pilosus* | 1 (<1%) |
| *Blaniulus guttulatus* | 1 (<1%) |
| *Proteroiulus fuscus* | 1 (<1%) |

### 3.2 Temporal relationships

Species appear to have clear seasonal patterns, with abundance minima and maxima in different seasons (Table 2); and different clades show different patterns. Earthworms have their abundance maxima in the winter months (December – February) and their abundance minima in the summer months (July and August) (Fig 1a). Woodlice and millipedes have similar patterns to each other but these differ markedly from the earthworm pattern, with their abundance maxima in the summer months (June – August) and their abundance maxima in the winter months (December to February) (Fig 1b). There appear to be no consistent yearly trends for any species, except *Dendrobaena attemsi*, which shows a plateauing increase in abundance over time, and *Porcellio scaber*, which shows a monotonic increase over time.

**Table 2.** Month of minimum and maximum abundances for each species of . The year response was tested against a null-hypothesis of no response at the p>0.05 level and that for a linear response at the p<0.05 level. H = estimated band width.

| Species | Minimum | Maximum | Year |
| --- | --- | --- | --- |
| *Chordeuma* sp | February | August | No change ($h>10$) |
| *Glomeris marginata* | December | July | No change ($h>10$) |
| *Trichoniscus pusillus* | December | June | No change ($h>10$) |
| *Porcellio scaber* | January | July | Rising, linear ($h>10$) |
| *Oniscus asellus* | January | July | No change ($h>10$) |
| *Philoscia muscorum* | January | July | No change ($h=4.57$) |
| *Aporrectodea caliginosa* | July | December | Lower in yrs 1 & 2 ($h = 2.53$) |
| *Allolobophora chlorotica* | August | February | No change ($h>10$) |
| *Dendrobaena attemsi* | August | February | Rises to plateau ($h=2.53$) |
| *Dendrobaena octaedra* | August | February | No change ($h>10$) |
| All earthworms | August | February | No change ($h>10$) |
| All woodlice | February | August | No change ($h>10$) |

**Figure 1:**
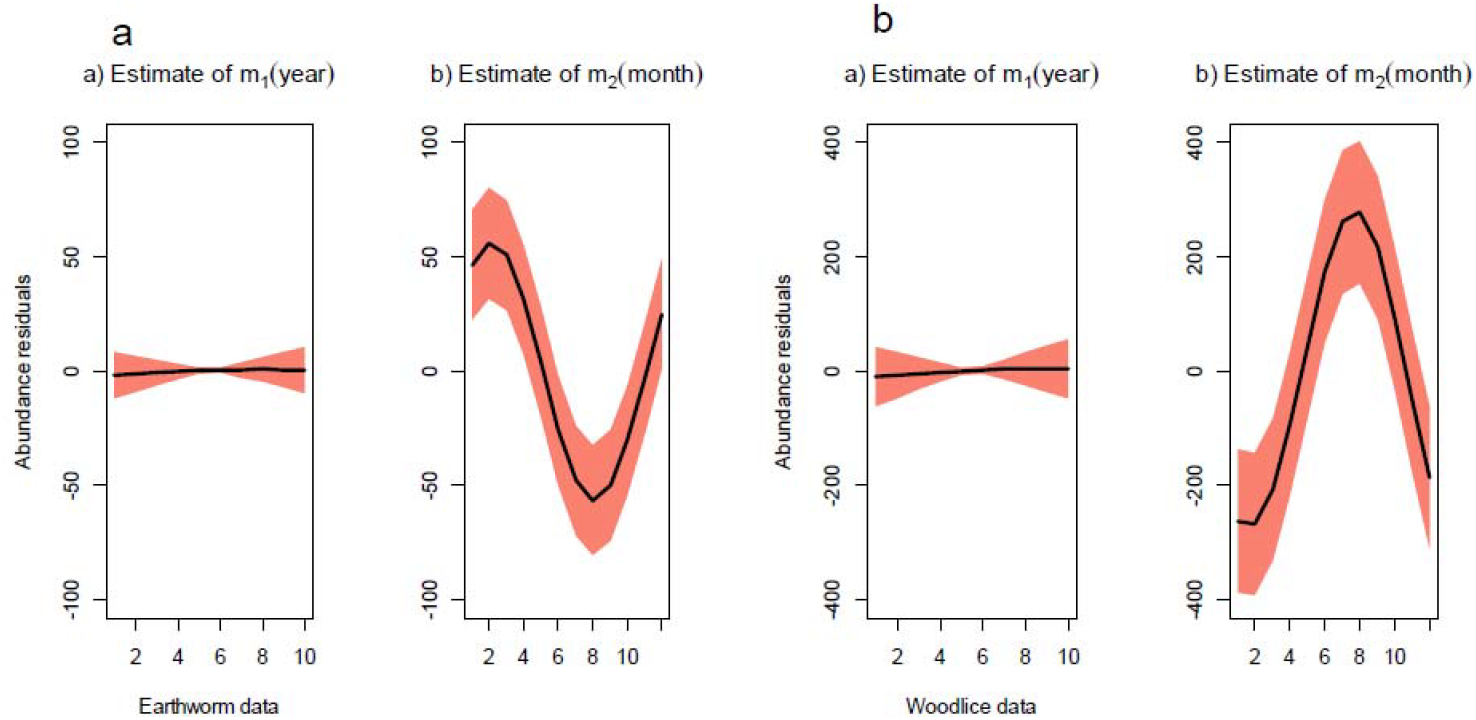
Non-parametric regressions for earthworm (a) and woodlice (b) abundance, with the variance partitioned into year and month.

### 3.3 Environmental (non-temporal) relationships

There is a strong (although not completely linear) negative relationship between soil temperature and soil moisture, such that whenever the soil temperature is high then soil moisture is low, and vice-a-versa (Fig 2). These conditions occur, of course, particularly in the extremes of the seasons (August and February).

**Figure 2:**
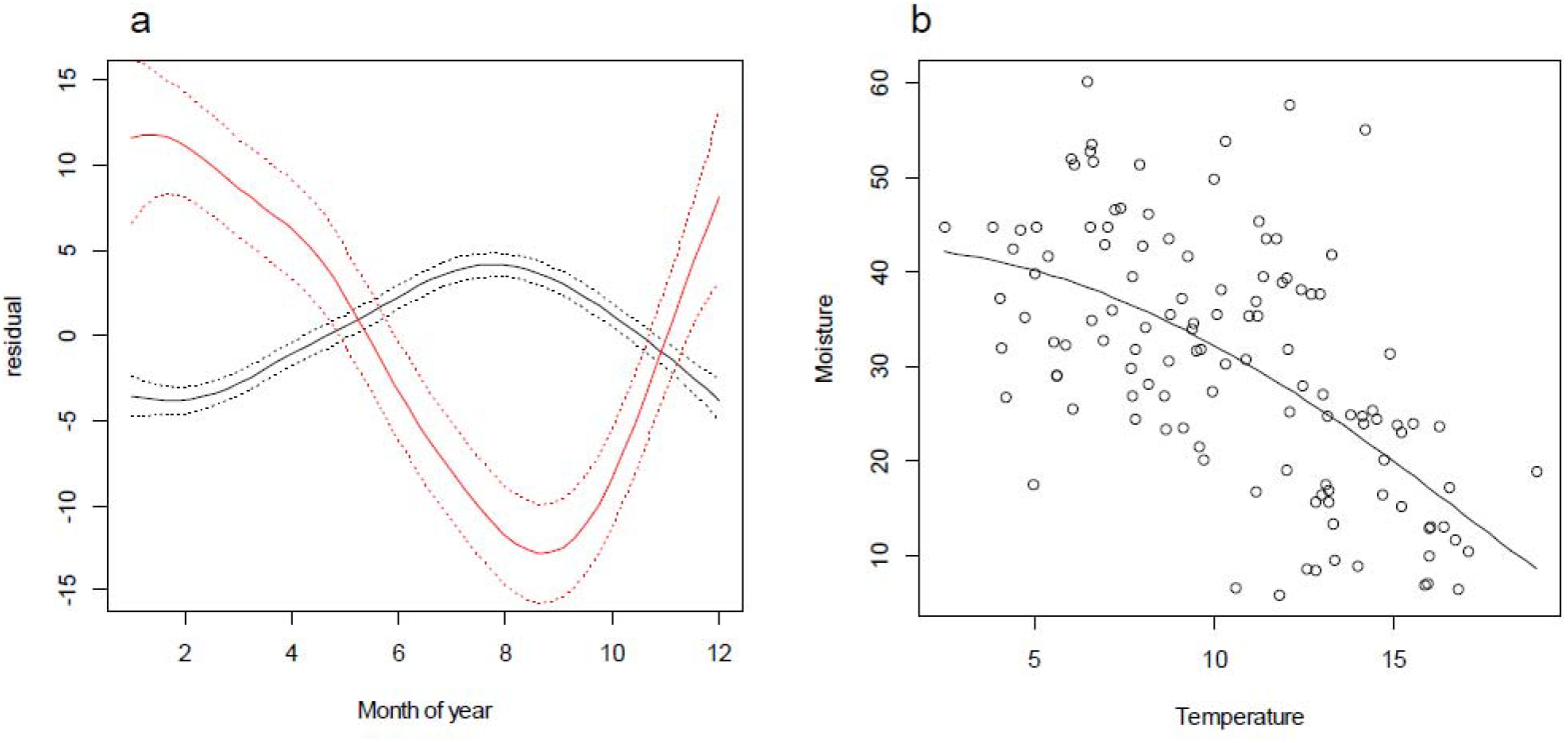
Relationship between temperature and moisture. (a) Non-parametric regression with variance partitioned into month. (b)non-parametric regression of temperature (black line) vs moisture (red line).

As a general pattern, then, earthworms and woodlice species have different and opposite responses to temperature and moisture, while millipedes show a similar pattern to woodlice. Earthworm abundance decreases with increasing temperature, whereas woodlice abundance increases with increasing temperature (Fig 3a).

**Figure 3:**
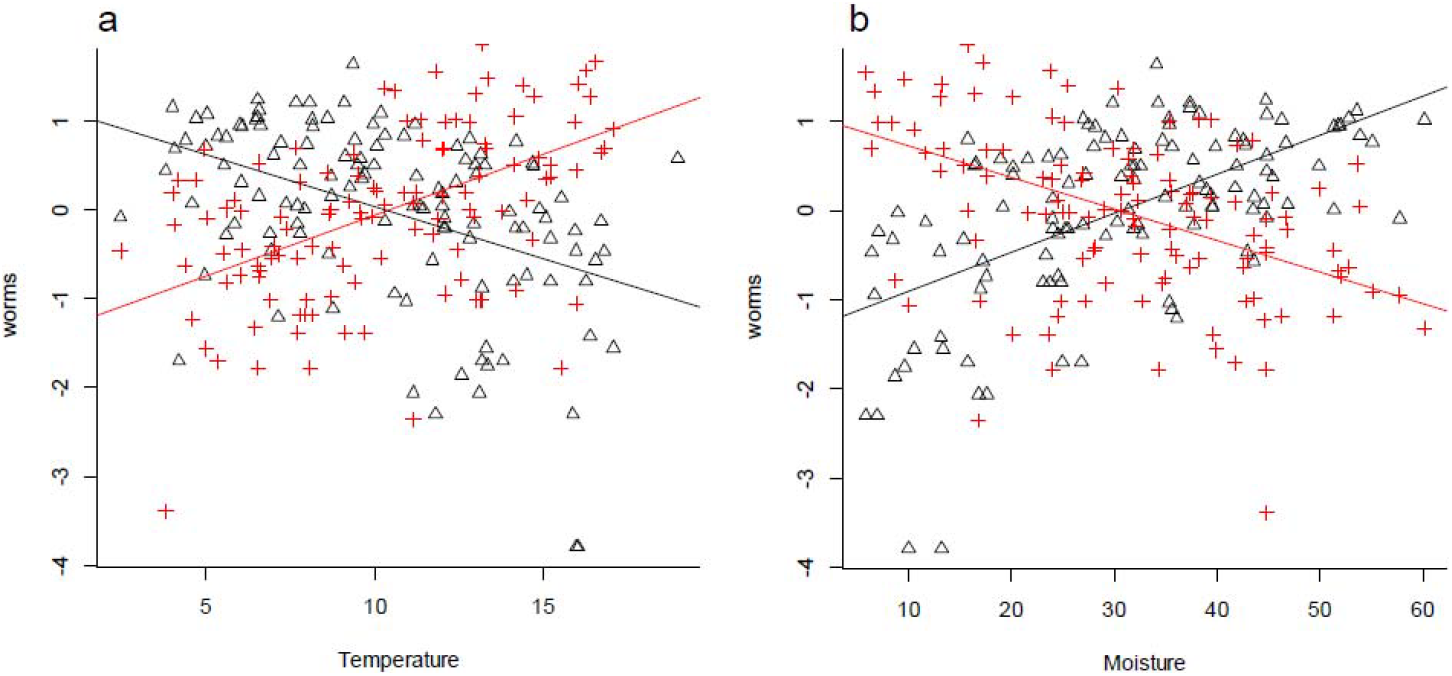
Non-parametric regressions of earthworms (black lines) and woodlice (redlines) log abundance against temperature (a) and moisture (b).

The opposite trend is true with respect to moisture, where earthworm abundance increases with increasing moisture and woodlice abundance decreases with increasing moisture (Fig 3b). This means that the highs and lows of both groups coincide with the seasonal temperature/moisture highs and low.

## 4. Discussion

We have established that woodlice and earthworms show contrasting patterns in their seasonal minima and maxima across the woodland plot. Woodlice are most abundant when earthworms are least abundant, and vice versa. This seems to be a result of the different environmental tolerances of the two groups and their different responses to the same environmental factors.

Our data reflects earlier findings that soil moisture is the most important environmental factor affecting earthworm abundance (e.g. Gerard, 1967; Auserwald et al., 1996; Cannavacciuolo et al., 1998). Soil moisture content is highest during the winter months when earthworms are at their most abundant and this because earthworms are extremely sensitive to desiccation. During drought conditions earthworms either aestivate (Edwards and Bohlen, 1996) or survive as cocoons (Petersen et al., 2008). Few adults can survive low soil moisture conditions, so when soil moisture is lowest during the summer months, earthworm abundance and activity is lowest. Earthworm abundance is also inversely correlated with temperature, but earthworms appear to be responding to the interaction of temperature and moisture (Gerard, 1967). When the soil temperature is lowest, soil moisture content is highest and vice versa. Frost conditions can severely affect earthworm abundance as only some species of earthworm (or their cocoons) are frost tolerant (e.g. *Dendrobaena octaedra* (Holmstrup and Loeschcke, 2003)). However, soil temperature did not drop below 2° C during the study period (minimum soil temperature, 2° C, maximum soil temperature, 19° C), and so earthworms were presumably able to remain active during all of the winter when soil moisture is generally at its highest.

In contrast, woodlice were most abundant in the summer months. This is probably because woodlice are known to respond to photoperiod (McQueen and Steel, 1980) and increase activity with increasing day length (Wieser, 1984). Daylight hours are greatest in summer when it is warmest, hence the positive correlation between abundance and increasing temperature. Day length is longest in June, shortly before the peak of activity for woodlice in this study. During the winter months, when the days are short, woodlice are therefore at their least active. Woodlice are well adapted to survive low moisture conditions, but are still limited by moisture (Wieser, 1984; Warburg et al., 1984) but possibly at soil moisture levels lower than found in this study.

Millipedes show similar patterns to woodlice, but their abundances are much lower and so they probably do not play as great a role in decomposition in Whitley Wood. Millipedes require calcium to build and repair their exoskeletons (Hopkin and Read, 1992). Soil pH in Whitley Wood is low (ca. 4.5) and calcium is likely to be limiting; the geology of the New Forest is made up of clays, gravels and sands (Melville and Freshney, 1982) and therefore there is likely to be only very small amounts of available calcium (Marschner, 1991). Consequently, millipede abundances are probably lower when compared with populations in woodlands growing on soils of higher pH over chalk or limestone (Lee, 2006).

This study shows that in our study woodland earthworms and woodlice (and, to a lesser extent, millipedes) have complementary seasonal patterns of density. Earthworms are at their most abundant when woodlice are at their least abundant, and vice versa. We propose two hypotheses that might explain the opposite seasonal patterns of these two important groups of macrofauna decomposers.

First, is true complementarity. The differences in peak abundance are due to abiotic rather than biotic factors; earthworms and woodlice do not appear to compete for leaf litter as their periods of peak abundance do not coincide. Earthworms and woodlice have similar ecological functions in the breaking down and incorporating of organic matter into the soil. It is therefore quite possible that this density complementarity means that rates of decomposition are maintained, at least approximately, throughout the year as a result of this complementary effect. However, the full extent of this ecological complementarity can only be assessed through proper litter decomposition experiments with separate exclusions of both groups.

A second possibility is resource partitioning by litter decomposition succession (Slade and Riutta, 2012; Riutta et al., 2012). In this hypothesis, competition between earthworms and woodlice results in seasonal partitioning of litter, with earthworms being more abundance during leaf abscission than woodlice. Within UK woodlands as a whole, leaf palatability of different species means that earthworms feed first on ash (very palatable) and then oak (less palatable), but they feed poorly on beech (unpalatable) (Quadros *et al.* 2014). Woodlice are unable to feed on ash as it has been consumed by earthworms, but can feed on oak and beech. Therefore, the leaf litter resource is partitioned according to leaf palatability. This could also be the result of past competition, through which selection pressure meant that the two groups partitioned the available litter resource seasonally. A combination of seasonal and litter succession would also explain patterns of seasonal abundance.

Sampling and extraction methods may have biased our results. Winkler bags are an effective method for sampling macrofauna from leaf litter, but there are large differences in the extraction efficiency for different taxa; Krell et al. (2005) showed that even after seven weeks of litter drying in Winkler bags, woodlice were still recovered. The moisture content of the litter also affects how rapidly invertebrates are extracted by Winkler bags, with wet litter having increased extraction times compared with dry litter (NHM Soil Biodiversity Group, unpublished data). It is likely that extractions from dry litter would yield greater numbers of woodlice than wet litter over a three day period. One would expect then that woodlice abundances would be greater in August and September when the litter is driest (see Eggleton et al., 2009 for a discussion of patterns in moisture content). However, woodlice abundance peaks in June and July when the litter is still relatively wet and therefore the litter moisture content bias cannot be an adequate full explanation for these results.

Increasingly warm and dry summers as a result of climate change (Murphy et al., 2009) could see reductions in earthworm numbers due to adults and/or cocoons not surviving the dry summer period (Holmstrup and Westh, 1995). This in turn could lead to a reduction in decomposition during the winter months due to lower earthworm densities. Some ecological compensation may occur with higher summer and winter woodlice abundances, with a corresponding increase in litter shredding. Woodlice incorporate litter into the soil (Hassall et al., 1987) but not to the same extent as earthworms, so litter incorporation may decrease. This in turn could lead to changes in soil structure accompanied by associated changes in soil fertility and pH (Riley et al., 2008). Dry soils particularly affect endogeic earthworms (Auserwald et al., 1996) and reductions in these species could affect soil structure and function. Reduced burrowing activity could lead to reduced porosity, aeration and drainage, fewer soil aggregates and less mixing of organic and mineral components of the soil (Lavelle, 1988).

This study illustrates the value of long term ecological data sets for uncovering ecologically meaningful seasonal patterns. We have found evidence for functional complementarity in two key decomposer groups that have seasonal peak abundances at different times of the year. This complementarity seems to be due to opposite responses to two major drivers: soil moisture and day length. This effect is likely to be widespread in lowland woodland in the UK and Europe, with only extreme temperatures (i.e. freezing soils or prolonged drought) and low pH limiting the distribution.

Functional complementarity has been documented widely in various different systems (e.g. Caradus et al., 1996; Stachowicz and Byrnes, 2004). Systems with high species diversity and functional complementarity are more resilient in the face of stress (e.g. invasion, changes in moisture and temperature) (Power, 2001). Function in ecosystems can be maintained by the complementary effects of functionally analogous species. It is increasingly important therefore that we understand the complementary effects of species in the face of environmental change, to understand which systems have in built resilience and which do not, and to focus efforts on restoring resilience to those system that have the greatest need.

## Acknowledgements

We thank the many people, especially Museum volunteers, too many to name individually, who have helped us with field work and sample sorting over the last ten years. Thanks to the staff at Tasty Pastries, Lyndhurst, for keeping us well fed for so long. Thank you to Simon Weymouth and Jayne Albery at the Forestry Commission, Lyndhurst, for granting access to Whitley Wood to conduct this study. This work has been supported financially by the Natural History Museum through its grant-in-aid funding. The authors declare that they have no conflict of interest with regards to this paper.

